# Stability of association between *Arabidopsis thaliana* and *Pseudomonas* pathogens over evolutionary time scales

**DOI:** 10.1101/241760

**Authors:** Talia L. Karasov, Juliana Almario, Claudia Friedemann, Wei Ding, Michael Giolai, Darren Heavens, Sonja Kersten, Derek S. Lundberg, Manuela Neumann, Julian Regalado, Richard A. Neher, Eric Kemen, Detlef Weigel

## Abstract

Crop disease outbreaks are often associated with clonal expansions of single pathogenic lineages. To determine whether similar boom-and-bust scenarios hold for wild plant pathogens, we carried out a multi-year multi-site survey of *Pseudomonas* in the natural host *Arabidopsis thaliana.* The most common *Pseudomonas* lineage corresponded to a pathogenic clade present in all sites. Sequencing of 1,524 *Pseudomonas* genomes revealed this lineage to have diversified approximately 300,000 years ago, containing dozens of genetically distinct pathogenic sublineages. These sublineages have expanded in parallel within the same populations and are differentiated both at the level of gene content and disease phenotype. Such coexistence of diverse sublineages indicates that in contrast to crop systems, no single strain has been able to overtake these *A. thaliana* populations in the recent past. Our results suggest that the selective pressures acting on a plant pathogen in wild hosts may be more complex than those in agricultural systems.

## Introduction

In agricultural and clinical settings, pathogenic colonizations are frequently associated with expansions of single or a few genetically identical microbial lineages (Butler et al., 2013; Cai et al., 2011; Kolmer, 2005; Park et al., 2015; Stukenbrock and McDonald, 2008; Yoshida et al., 2013). The conditions that lead to such epidemics-such as reduced host genetic diversity (Zhu et al., 2000), absence of competing microbial communities (Brown et al., 2013) or high transmission rates (Park et al., 2015)-are, however, by no means a universal feature of pathogenic infections. Instead, many, if not most, pathogens can colonize host populations that are both genetically diverse and that can accommodate a diversity of other microbes (Barrett et al., 2009; Falkinham et al., 2015; Woolhouse et al., 2001).

Factors that drive pathogen success in such more complex situations are less well understood than for clonal epidemics. For example, if a pathogen species persists at high numbers in non-host environments, does each host become infected by a different pathogen strain? Or does a multitude of genetically distinct pathogens infect each host? And do different colonizing strains use disparate mechanisms to become established even within genetically similar host individuals? The answers to these questions inform on how (and if) a host population can evolve partial or even complete pathogen resistance (Anderson and May, 1982; Barrett et al., 2009; Karasov et al., 2014a; Laine et al., 2011). Several studies over the past 20 years have attempted to infer the distributions of non-epidemic pathogens in both host and non-host environments (Falkinham et al., 2015; Wiehlmann et al., 2007). These studies, which have observed a range of different patterns, are unfortunately often limited to the historic strains that are available, and the conclusions vary for different collections, even of the same pathogen species (Pirnay et al., 2009).

Questions of pathogen epidemiology are of particular relevance when considering the genus *Pseudomonas,* which includes pathogens and commensals of both animals and plants (Baltrus et al., 2017) and is among the most abundant genera in plant leaf tissue. This genus belongs to the Gram-negative gammaproteobacteria, with well over a hundred recognized species (Gomila et al., 2015). The three taxa most commonly found on plants are *P. syringae* and *P. viridiflava* in the *P. syringae* complex (Bartoli et al., 2014) and *P. fluorescens* (Garrido-Sanz et al., 2016). The abundance of *Pseudomonas* can have a large impact on plant fitness (Balestra et al., 2009; Gao et al., 2009; Yunis et al., 1980), and several putatively host-adapted lineages of this genus (Baltrus et al., 2011, 2012) can cause agricultural disease epidemics. Despite the damage they can do to plants, *Pseudomonas* pathogens are not obligatory biotrophs: surveys of *Pseudomonas* in environmental and nonhost habitats have revealed distribution patterns typical for opportunistic microbes (Bartoli et al., 2014; Morris et al., 2008, 2010), with genetically divergent lineages not uncommonly found in the same host populations (Barrett et al., 2011; Karasov et al., 2017; Kniskern et al., 2011).

Motivated by wanting to understand how the distribution of a common plant pathogen differs between agricultural and non-agricultural situations, we have begun to elucidate the epidemiology of *Pseudomonas* strains within and between populations of a non-agricultural host. *Arabidopsis thaliana* is a globally distributed wild plant capable of colonizing poor substrates as well as fertilised soils (Weigel, 2012). *Arabidopsis thaliana* populations across the globe are hosts to *Pseudomonas,* and several of the most abundant *Pseudomonas* strains are pathogenic on *A. thaliana,* even though they are likely not specialized on this species as a host (Barrett et al., 2011; Bodenhausen et al., 2014; Cai et al., 2011; Jakob et al., 2002, 2007; Kniskern et al., 2011).

Here we report a broad-scale survey of *Pseudomonas* operational taxonomical units (OTUs) based on 16S rDNA sequences in six *A. thaliana* populations from South-Western Germany, over six seasons. Through this survey we first identified a single OTU that was consistently dominating in individual plants, across populations and across seasons. Through subsequent isolation and sequencing of the genomes of 1,524 *Pseudomonas* isolates we uncovered extensive diversity within this pathogenic OTU, diversity that is much older than *A. thaliana* in this area. Taken together, this makes for a colonization pattern that differs substantially from what is typically observed for crop pathogens. The observation of a single dominant and temporally persistent *Pseudomonas* lineage in several host populations is at first glance reminiscent of successful pathogens in agricultural systems. However, in stark contrast to many crop pathogens, this *Pseudomonas* pathogen can apparently persist as a diverse metapopulation over long periods, without a single sublineage becoming dominant.

## Results

### Dozens of *Pseudomonas* OTUs persist in *A. thaliana* populations

*Pseudomonas* bacteria are abundant in *A. thaliana* populations from South-Western Germany (Agler et al., 2016), but whether the same lineages are found in these different populations and whether the abundant lineages are pathogenic was not known. To obtain a first understanding of *Pseudomonas* diversity on *A. thaliana,* we surveyed 6S rDNA diversity across six host populations in spring and fall of three consecutive years. We sampled both the epiphytic and endophytic microbiome of rosettes and sequenced the v3-v4 region of 16S rDNA (Fig. 1a, Fig. S1a, Table S1). As expected, *Pseudomonas* was common, occurring in 92% of samples (97% of the epiphytic and 88% of the endophytic samples) and representing on average 3% of the total bacterial community. The genus was found at similar densities inside and on the surface of leaves (ANOVA, P>0.05) (Fig. S1 b), indicating no preferential colonization of either niche. While we did not detect an effect of sampling time on relative abundance (ANOVA, P>0.05), abundance varied across sites (ANOVA, R^2^=12.8%, *P*=10^-10^; Fig. S1c), suggesting that certain site-specific characteristics may be particularly conducive to *Pseudomonas* proliferation.

**Fig. 1.**
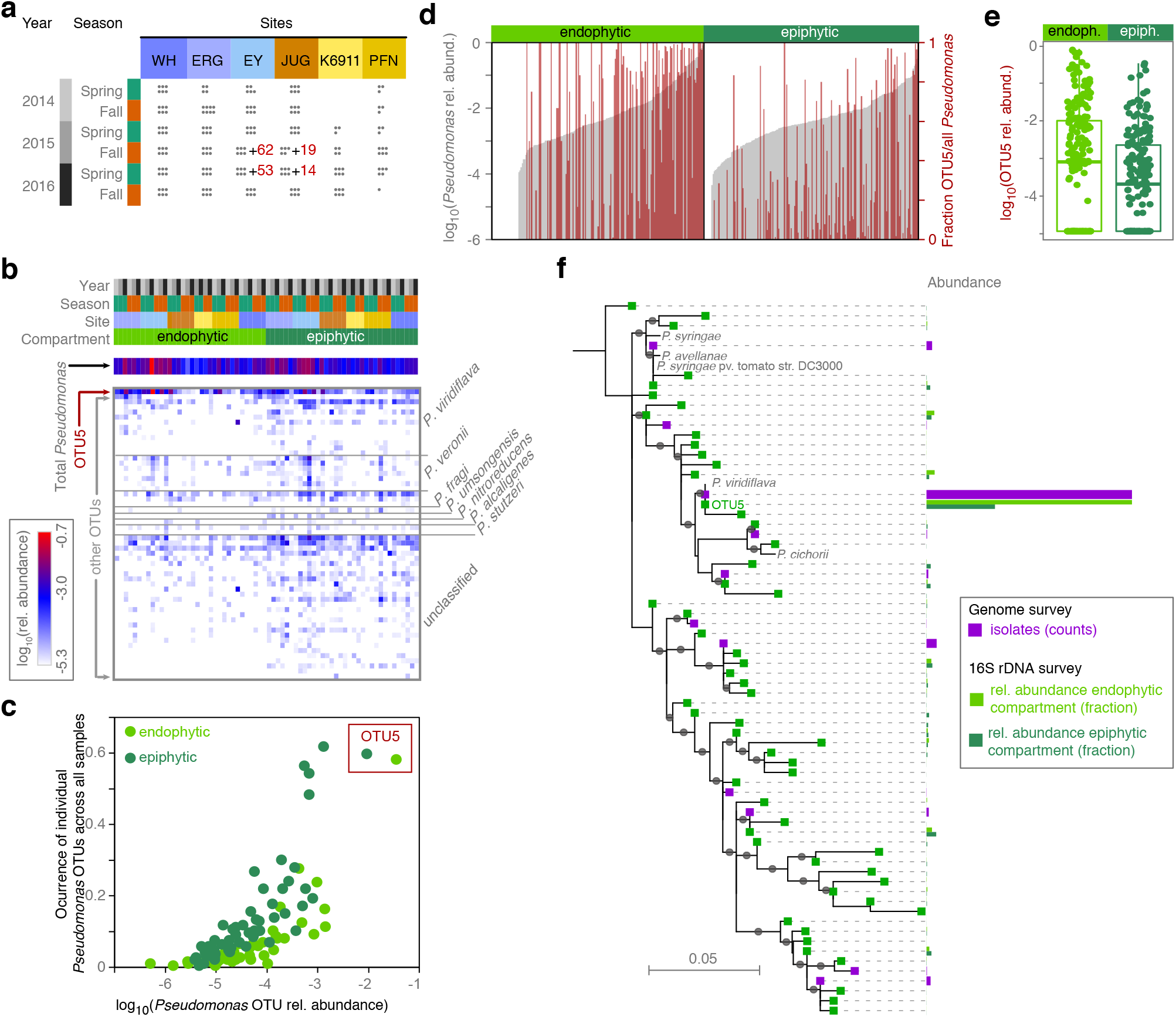
Natural *Pseudomonas* populations in *A. thaliana* leaves are dominated by the OTU5 lineage. (a) Overview of 16S rDNA survey of epi-and endophytic compartments of *A. thaliana* plants (dots indicate sampled plants). Numbers of individuals from which *Pseudomonas* isolates were cultured and metagenome analysis was performed in parallel are indicated in red. (b) Heat map of relative abundance of 56 *Pseudomonas* OTUs in the 16S rDNA survey. Color key to samples on top according to panel (a). *Pseudomonas* species assignments on the right. *P. veronii, P. fragi* and *P. umsongensis* belong to the *P. fluorescence* complex, *P. nitroreducens* and *P. alcaligenes* to the *P. aeruginosa* complex. (c) Correlation between occurrence across all samples and average relative abundance within samples of the 56 *Pseudomonas* OTUs in the endo-and epiphytic compartments. (d) *Pseudomonas* abundance (grey bars) and percentage of *Pseudomonas* reads belonging to OTU5 (red bars), in the endo-and epiphytic compartments. (e) OTU5 is significantly more abundant in the endophytic compartment (Wilcoxon test, *P* = 0.02). (f) Maximum-likelihood phylogenetic tree illustrating the similarity between amplicon sequencing derived and isolation derived *Pseudomonas* OTUs defined by distance clustering at 99% sequence identity of the v3-v4 regions of the 16S rDNA. For isolates, exact 16S rDNA sequences were used, for amplicon sequencing OTU the most common representative sequence was used. Grey dots on branches indicate bootstrap values >0.7. Color bars represent the relative abundance or the number of isolates. The most abundant *Pseudomonas* OTU in both the endophytic and epiphytic compartments, OTU5, was identical in sequence to the most abundant sequence observed among isolates and to a *P. viridiflava* reference genome (NCBI AY597278.1/AY597280.1). See also Fig. S1 and S2.

By clustering of *Pseudomonas* 16S rDNA reads at 99% sequence similarity, we could distinguish 56 OTUs (Fig. 1b). The 99% threshold for distance-based clustering of reads resulted in OTU patterns more congruent with a whole genome-phylogeny than the more widely used 97% sequence similarity (Fig. S2). While half of the *Pseudomonas* OTUs could not be classified at the species level, 13 were classified as *P. viridiflava,* which belongs to the *P. syringae* complex, including the most abundant OTU, OTU5. The other classifiable OTUs belonged to the *P. fluorescens, P. aeruginosa* and the *P. stutzeri* species complexes (Fig. 1b).

To understand the factors shaping *Pseudomonas* assemblages, we studied variation in OTU presence and relative abundances as an indication of *Pseudomonas* population structure. Permutational multivariate analysis of variance (PerMANOVA on Bray-Curtis distances, *P*<0.05) indicated that differences between host individuals were associated primarily with interactions between site, leaf niche and sampling time (20.0% explained variance), with a smaller percentage associated with each factor independently such as site (4.0% explained variance), leaf niche (2.3%) or sampling time (2.7%). An important difference between leaf niches was that endophytic *Pseudomonas* populations were 2.6 times less diverse than epiphytic populations (Wilcoxon test, P< 10^-16^) (Fig. S1 d), pointing to selective bottlenecks inside the leaf being stronger.

### A single lineage dominates *Pseudomonas* populations in *A. thaliana* leaves

*Pseudomonas viridiflava* OTU5 was overall the most common *Pseudomonas* OTU across samples (Fig. 1b), occurring in 59% of epiphytic and 58% of endophytic samples. Across all samples, OTU5 accounted for almost half of reads in the endophytic compartment (48%, range 0-99.9% in each sample), and it was the most abundant endophytic *Pseudomonas* OTU in 52% of samples (Fig. 1c). The dominance of OTU5 was less pronounced in the epiphytic samples, where it averaged 23% of all reads (range 0-99.9%), being the most abundant OTU in only 23% of samples. This observation indicates an enrichment of this OTU in the endophytic over the epiphytic compartment (Wilcoxon test *P* = 0.02, paired Wilcoxon test *P* = 7.4x10^-9^) (Fig. 1d). In conjunction with the reduced *Pseudomonas* diversity in the endophytic compartment, this is evidence for OTU5 members being particularly successful endophytic colonizers of *A. thaliana.*

16S rDNA amplicon reads reveal the relative abundance of microbes, but they do not inform on the absolute abundance of microbial cells in a plant, what we term the ‘microbial load’. The latter is perhaps a more informative readout of the selective pressure exerted by a single microbial taxon than the relative abundance of a taxon among all microbes. A pathogen might dominate the microbiota, but unless it reaches a certain abundance, there might not be a marked decrease in host fitness (Duchmann et al., 1995; Schneider and Ayres, 2008; Vaughn et al., 2000). The importance of absolute microbial load has recently come into focus of human gut microbiome analyses as well (Vandeputte et al., 2017).

To determine whether *Pseudomonas* diversity was related to overall microbial load, we used metagenome shotgun sequencing to quantify total microbial colonization. We returned to two of the previously sampled populations (Fig. 1a; Fig. S1 a), collected and extracted genomic DNA from entire, washed leaf rosettes, and performed whole-genome shotgun sequencing on 192 plants. The same DNA was also used for 16S rDNA amplicon analysis to call OTUs. We mapped Illumina sequencing reads against all bacterial genomes in GenBank and against the *A. thaliana* reference genome, and determined the ratio of bacterial to plant reads. We calculated the correlation of microbial load, which varied substantially across the 192 plants (Fig. 2a), with each of the 6,715 OTUs detected in at least one sample. Because OTUs were called on 16S rDNA amplicon sequences, but microbial load was assessed on metagenomic reads, the two assays provided independent measurements of relative and absolute microbe abundance. Among all OTUs, OTU5 was the second most highly correlated with total microbial load (Fig. 2b,c; Pearson correlation coefficient R=0.41, q-value=7x10^-5^), indicating that the strains represented by OTU5 are not only the most common *Pseudomonas* strains in these plants, but also that they are either major drivers or beneficiaries of microbial infection in these plants.

**Fig. 2.**
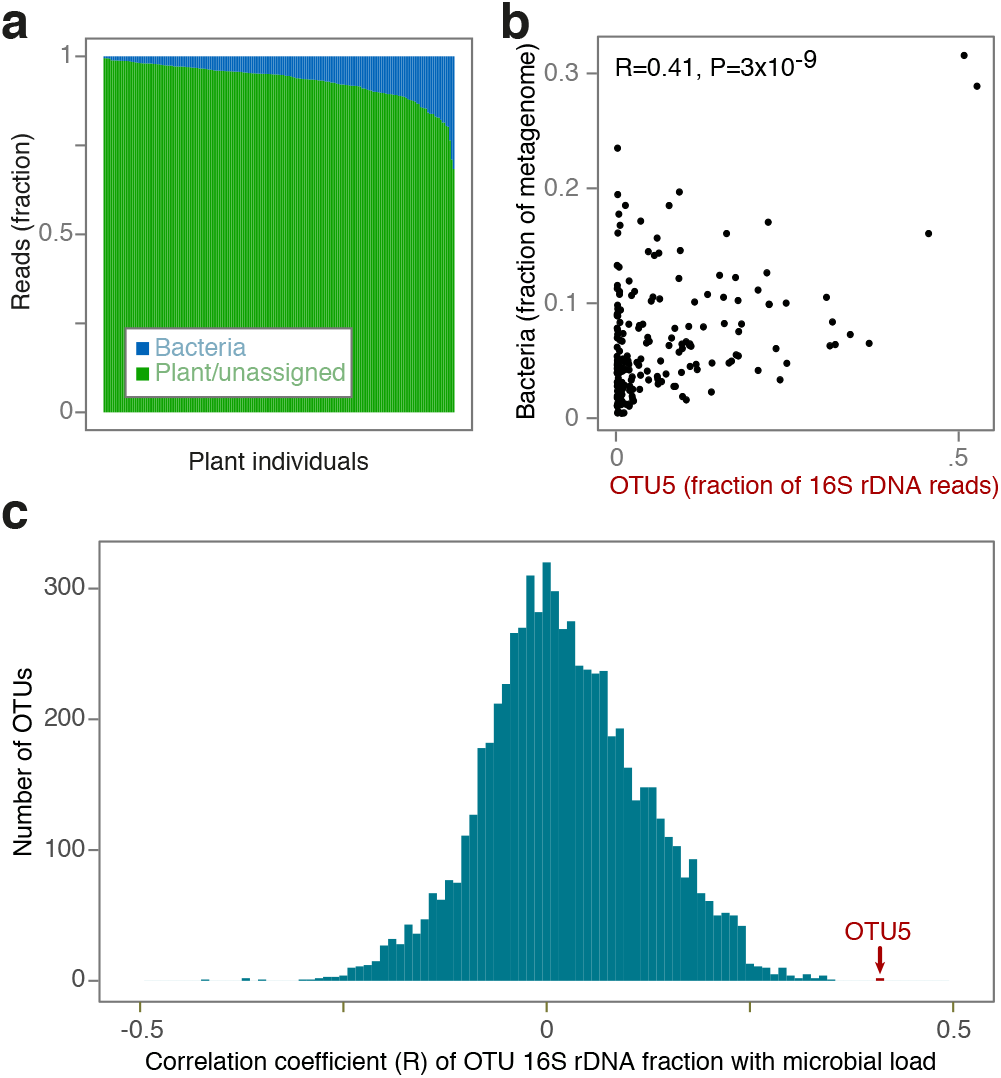
The most abundant OTU, OTU5, is correlated with microbial load. (a) Bacterial and plant fraction of metagenome shotgun sequencing reads in 192 plants. (b) Correlation between fraction of bacterial reads in metagenome data and relative abundance of OTU5 in 16S rDNA amplicons from the same 192 samples. (c) Distribution of Pearson correlation coefficients between microbial loads as inferred from fraction of bacterial reads and OTU abundances (as shown for OTU5 in panel (b)). The correlation coefficient for OTU5 abundance is the second highest correlation among 6,715 OTUs detected across all samples. See also Fig. S2.!!

### OTU5 comprises many genetically distinct strains

While OTU classification based on 16S rDNA and metagenomic assignment can be indicative of the genus or species-level identity of a microbe, genetically and phenotypically diverse strains of a genus will often be clustered together as a single OTU (Moeller et al., 2016). To discern genetic differentiation within OTU5, we therefore wanted to compare the complete genomes of OTU5 strains. From the same plants in which we had analyzed the metagenomes, we cultured and isolated between 1 and 34 *Pseudomonas* colonies (mean= 11 per plant, median=12). We then sequenced and assembled *de novo* the full genomes of 1,611 *Pseudomonas* isolates (assembly pipeline and statistics in Fig. S3). Eighty-seven genomes with poor coverage, abnormal assembly characteristics or incoherent genome-wide sequence divergence were removed from further analysis. The remaining 1,524 genomes were 99.5% complete, as estimated with published methods (Simão et al., 2015), containing on average 5,347 predicted genes (standard deviation 284). Extraction of 16S rDNA sequences from the whole genome assemblies demonstrated that the vast majority of all isolates, 1,355, belonged to the OTU5 lineage, as defined previously by amplicon sequencing.

Maximum-likelihood (ML) whole-genome phylogenies (Ding et al., 2018) were constructed from the concatenation of 807 genes that classified as the aligned soft core genome of our *Pseudomonas* collection. Because bacteria undergo homologous recombination, the branch lengths of the ML whole-genome tree may not properly reflect the branch lengths of vertically inherited genes, but the overall topology is expected to remain consistent (Hedge and Wilson, 2014). The 1,524-genome phylogeny revealed hundreds of isolates that were nearly or completely identical across the core genome to at least one other isolate. Using a similarity cutoff of 99.9967% sequence identity (corresponding to a SNP approximately every 30,000 bp across the core genome based on distance in the ML tree), the 1,524 isolates collapsed into 189 distinct *Pseudomonas* strains (Fig. 3a). In the whole-genome tree, 1,355 OTU5 isolates, comprising 107 distinct strains, formed a single monophyletic clade. One genome (p8.A2) in this clade differed in its 16S rDNA taxonomical assignment, but was later found to be likely a mixture of two genomes. In support of the 16S rDNA placement of OTU5 within the *Pseudomonas* genus, the OTU5 clade is most closely related to *P. viridiflava* and *P. syringae* strains (Fig. 1f). Genetic differences between the identified strains were distributed through the genome, indicating that divergence between strains was not solely the result of a few importation events of divergent horizontally transferred material (Fig. 4a).

**Fig. 3.**
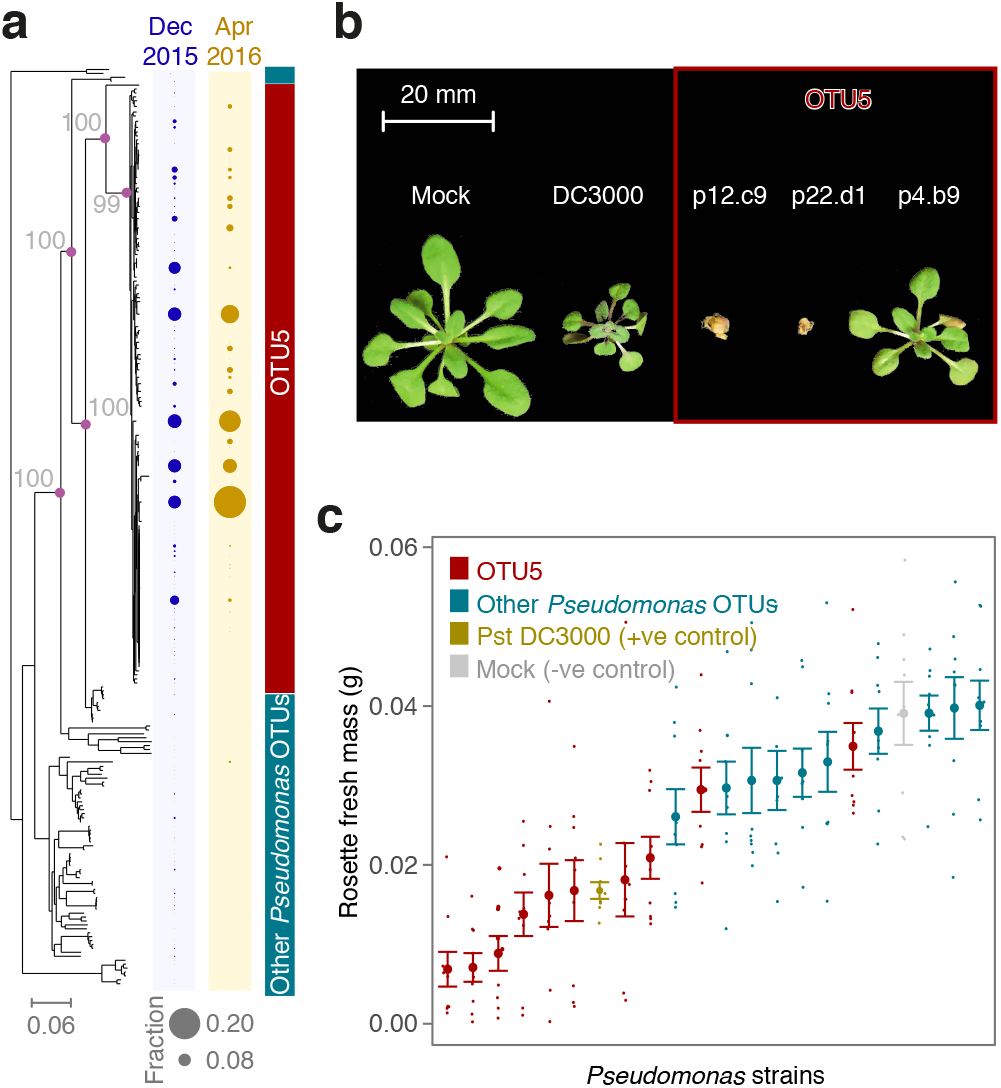
OTU5 is composed of multiple expanding lineages that are pathogenic. (a) ML whole-genome phylogeny and abundance of strains in Eyach, Germany, in December 2015 and April 2016. Diameters of circles on the right indicate relative abundance across all isolates from that season. Purple circles at nodes relevant for OTU5 classification and grey numbers indicate support with 100 bootstrap trials. (b) Examples of OTU5 strains that can reduce growth and even cause obvious disease symptoms in gnotobiotic hosts. (c) Quantification of effect of drip infection on growth of plants. Pst DC3000 was used as positive control. The negative control did not contain bacteria. See also Fig. S2 and S5.

**Fig. 4.**
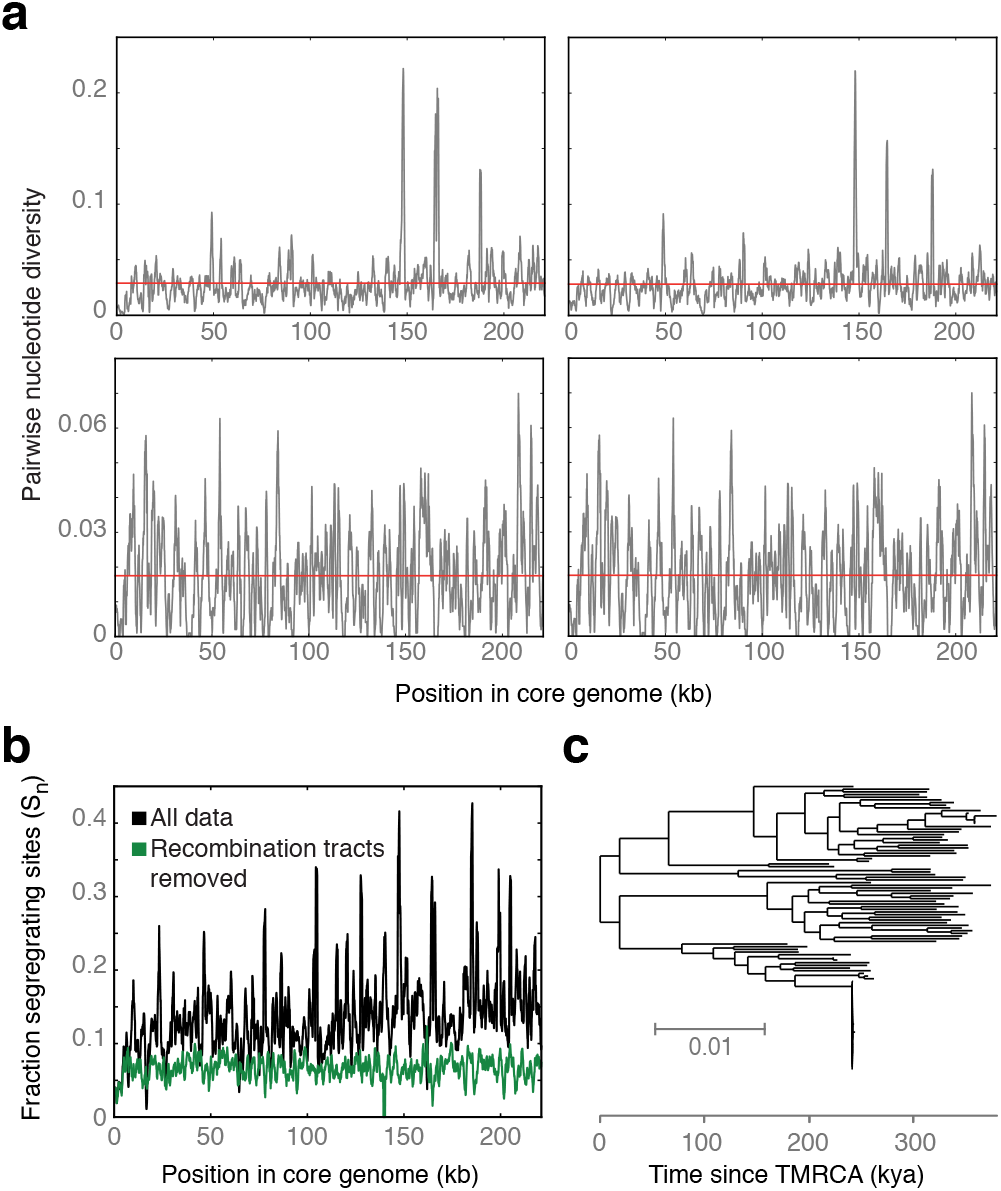
Genome-wide divergence and dating of OTU5 strains. (a) Pairwise nucleotide diversity in 1000 bp sliding windows. One randomly chosen OTU5 reference strain was separately compared with four different other OTU5 strains. (b) Genome-wide distribution of Segregating Sites (S_n_) in OTU5, calculated in 1000 bp sliding windows. Putative recombination tracts were removed from the core genome alignment to calculate the coalescence of OTU5. This removal reduced the fraction of segregating sites by half (0.14 vs. 0.07). (c) The TMRCA of 107 isolates representing the 107 identified OTU5 strains as calculated using a substitution rate estimated in ref. (McCann et al., 2017). See also Fig. S3.

Comparing the position of strains on the phylogeny and their provenance identified several strains that were not only frequent colonizers across plants, but also persistent colonizers over time, each isolated in at least two consecutive seasons (a). Six OTU5 strains accounted for 51% of sequenced isolates, each with an overall frequency of between 4-10%, with several found in over 20% of plants. In contrast, no strain outside of OTU5 exceeded an overall frequency of 5%. Generally, non-OTU5 strains were much less likely to be represented by multiple isolates and were very rarely observed in both seasons sampled.

### OTU5 primarily comprises pathogenic strains, but with distinct phenotypes

The *P. syringae/P. viridiflava* complex, to which OTU5 belongs, contains many well-known plant pathogens-although not all *P. syringae* complex strains are pathogenic, with some lacking the canonical machinery required for virulence (Barrett et al., 2011; Clarke et al., 2010). Because some infection characteristics are determined by the presence of a single or few genes, even closely related strains of the same species can cause diverse types of disease (Barrett et al., 2011; Nowell et al., 2016). Given the known phenotypic variability within and between *Pseudomonas* species, 16S rDNA sequences alone did not inform on the pathogenic potential of the OTU5 strains.

To determine directly the virulence-which we define here as the ability to cause disease-of diverse OTU5 isolates, we drip-inoculated 26 of them on seedlings of Eyach 15-2, an *A. thaliana* genotype common at one of our sampling sites in Southwestern Germany (Bomblies et al., 2010) (Fig. 3b-c, Fig. 4a; Fig. S5). Twenty-five of the 26 tested OTU5 strains reduced plant growth significantly in comparison to uninfected plants, but only two of ten randomly-chosen non-OTU5 *Pseudomonas* strains did so (ANOVA, P<0.05). The tomato pathogen *P. syringae* pv. tomato (Pst) DC3000, which is known to be highly virulent on *A. thaliana* (Velásquez et al., 2017), reduced growth to a similar extent as several OTU5 strains, with some OTU5 strains being even more virulent and killing seedlings outright (Fig. 3b,c). Treatment of seedlings with boiled, dead bacteria did not reduce plant growth for any of the five isolates tested (ANOVA, P>0.25 for all), indicating that the reduction in plant growth was not due to run-away immunity triggered by the initial inoculation, but was indeed caused by proliferation of living bacteria. From the clear phenotypic stratification of strains we conclude that the majority of OTU5 strains is virulent. These experimental results in conjunction with the observed correlation of OTU5 with microbial load in the field, established with metagenomic methods, both point to OTU5 as being responsible for some of the most persistent bacterial pathogen pressures in the sampled *A. thaliana* populations.

### Strains within OTU5 diverged over 300,000 years ago

Several surveys of crop pathogen epidemics have indicated that few, if not single strains frequently drive such outbreaks, with the dominant strains often changing over the course of a few years or decades (Cai et al., 2011; Kolmer, 2005; McCann et al., 2017; Stukenbrock and McDonald, 2008; Yoshida et al., 2013, 2014). An example of these dynamics has been well illustrated by Cai and colleagues (Cai et al., 2011) who followed the expansion of *P. syringae* strains in agricultural tomato populations during the 20^th^ century, finding that at nearly all time points only one or two strains were present at high frequency. Isolates from the lineage that was most abundant over the last sixty years-to which today over 90% of assayed isolates belong-differed at only a few dozen SNPs throughout the genome, indicative of a common ancestor as recently as just a few decades ago.

Our comparative analysis of the OTU5 lineage from *A. thaliana* had shown that OTU5 isolates were much more diverse, with over 10% of positions (27,217/221,628 bp) in the OTU5 core genome being polymorphic. However, the age of diversification cannot be inferred directly from a concatenated whole genome tree (Hedge and Wilson, 2014) because recombination events with horizontally transferred DNA can increase the sequence divergence between strains, thereby elongating branches and inflating estimates of the time to the most recent common ancestor (TMRCA). To prevent the overestimation of the TMRCA of OTU5, it was necessary to correct for the effects of recombination. Such correction can lead instead to underestimation of branch lengths (Hedge and Wilson, 2014); we found this acceptable, because our goal was to assess a minimum lower bound for TMRCA for the strains of interest. We removed 7,646 recombination tracts from the whole-genome alignments after having inferred recombination sites in the core genome using ClonalFrameML (Didelot and Wilson, 2015) (Fig. 4b). Removal of recombination tracts reduced the number of segregating sites by approximately 50%. As expected, the remaining polymorphic sites were distributed more evenly throughout the genome (Fig. 4b). For inference of neutral coalescence, it is ideal to consider substitutions at fourfold degenerate sites. However, limiting the analysis to fourfold degenerate sites after subsetting to a strict core genome and removal of putative recombination sites left too few segregating sites to make robust phylogenetic partitions. Hence, we performed subsequent calculations on all non-recombined sites.

McCann and colleagues (McCann et al., 2017) have used temporal collections of a clonally spreading kiwi pathogen to estimate the rate of substitution in a *Pseudomonas* lineage related to OTU5. Using their point estimate of 8.7x10^-8^ substitutions per site per year, we estimated the TMRCA of the 107 OTU5 strains. The ML-tree of OTU5 strains with recombination events removed contained a median mid-point-root to tip distance of 0.026 (standard deviation=0.004). With the substitution rate estimated by McCann and colleagues (McCann et al., 2017) this corresponds to a TMRCA estimate of 300,000 years (standard deviation of root-to-tip distances=46,000 years) (Fig. 4c).

Note that this is likely an underestimate of the TMRCA, due to removal of ancient homoplasies identified by ClonalframeML (Hedge and Wilson, 2014). Furthermore, the substitution rate estimate from McCann and colleagues (McCann et al., 2017) is likely higher than the long-term substitution rate relevant to OTU5 (Exposito-Alonso et al., 2018; Kryazhimskiy and Plotkin, 2008; Rocha et al., 2006). Nevertheless, from this data we can conclude that strains of OTU5 likely diverged from one another approximately 300,000 years ago, pre-dating the recolonization of Europe by *A. thaliana* from Southern refugia after the Last Glacial Maximum (1001 Genomes Consortium, 2016).

### Individual pathogenic strains often dominate *in planta*

Since multiple isolates (between one and 34, with a median of 12) had been sequenced from most sampled plants, we could assess the frequency of specific strains not only across the entire population, but also within each individual host. Most plants (73.3%) were colonized by multiple strains. While similar numbers of distinct strains within and outside of OTU5 were represented in our population level survey (Fig. 3a), non-OTU5 strains tended to be found at low frequencies in collections from individual plants (a,b). Of all OTUs, only OTU5 strains, most of which are pathogenic, were likely to partially or completely dominate, i.e., reach frequencies above 50%, within a single plant (Fig. 5a,b).

**Fig. 5.**
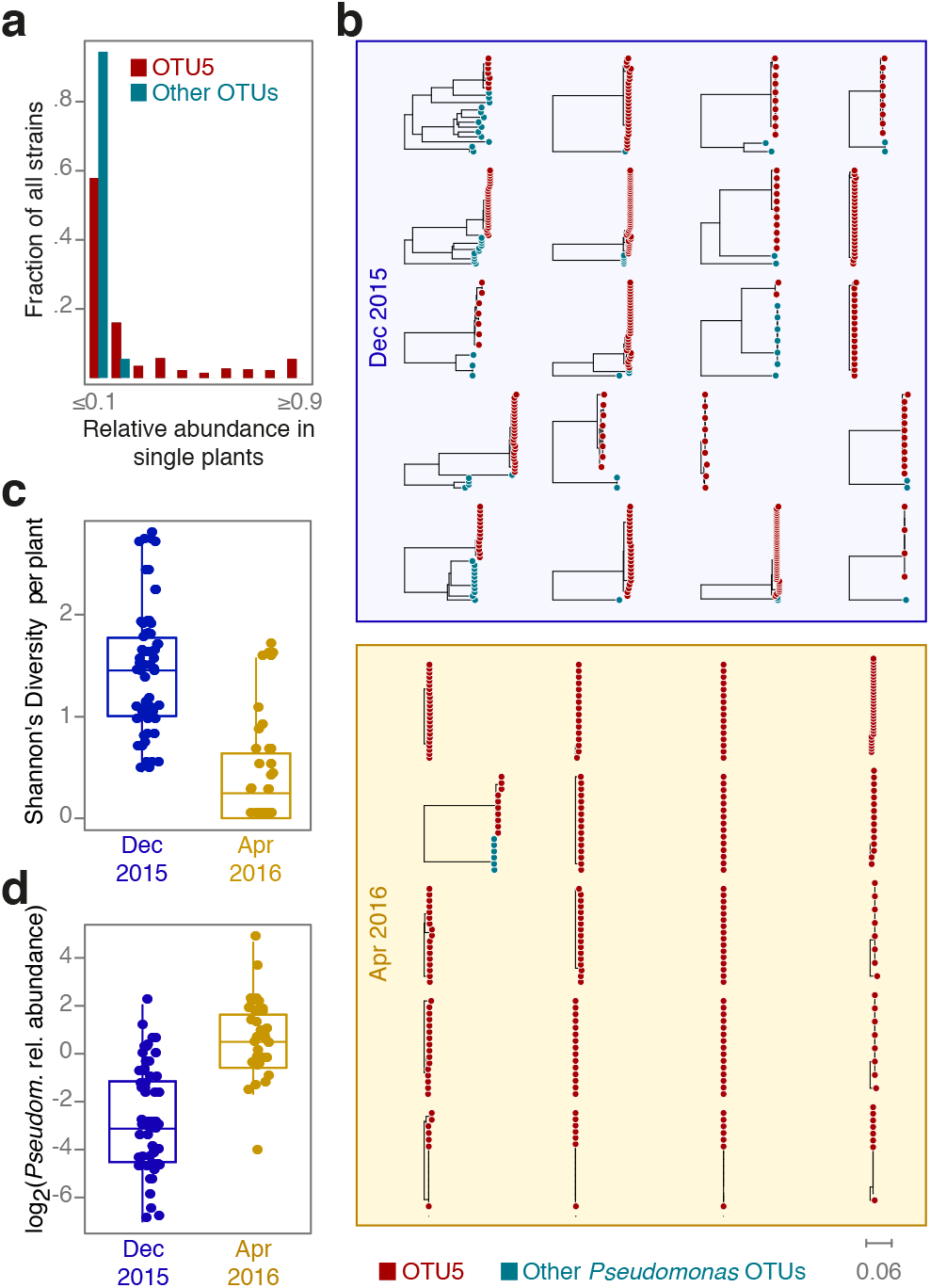
Different OTU5 strains expand clonally within different plants. (a) Distribution of relative OTU5 and non-OTU5 strain abundances in single plants. (b) Phylogenetic trees of isolates collected from individual plants. (c) Strain diversity as function of season. (d) *Pseudomonas* load as function of season. For both (c, d), seasons are significantly different (Student's t-test, P=l.32x10^-15^). Box-plots show median, first and third quartiles. Related to Fig. S4.

Strain diversity per host individual not only differed between clades, but also between seasons, with the distribution of strain frequency per plant changing over time. We measured the Shannon Index H’ (Hill, 1973) to compare strain diversity per plant across the two seasons in which we had sampled isolates. While the fall cohort tended to have been colonized by several strains simultaneously, plants in spring were characterized by reduced strain diversity (Fig. 5c) (Student's t-test, P=l.3x10^15^). One possible explanation for this change in strain frequencies is a local spring bloom of OTU5 populations. Plants sampled in spring carried a significantly higher absolute *Pseudomonas* load (Fig. 5d) (Student's t-test, P=l.0x10^-5^), consistent with spring conditions favoring local OTU5 proliferation.

### Gene content differentiation of the pathogen clade OTU5

The abundance of OTU5 as well as its enrichment in the endophytic over the epiphytic compartment indicated that this lineage colonizes *A. thaliana* more effectively than do related OTUs. Whether this success is the result of expansion in the plant, or host filtering of colonizers (Costello et al., 2012), is unclear, and we were curious what endows OTU5 strains with capacity to apparently outcompete other *Pseudomonas* lineages and to dominate in populations and in individual plants. To begin to answer this question, we sought to investigate a potential common genetic basis. To this end, we assessed the distribution of ortholog groups across the genomes of all *Pseudomonas* isolates including OTU5 using panX (Ding et al., 2018) (Fig. 6). From a presence-absence matrix in the pan-genome analysis one can immediately distinguish OTU5 lineages from non-OTU5 lineages. Nine hundred and fourteen genes are conserved (>90% of genomes) within OTU5, but are much more rarely found outside this OTU, in fewer than 10% of non-OTU5 strains. Most of the conserved genes, 59%, encode proteins without known function (“hypothetical proteins”).

**Fig. 6.**
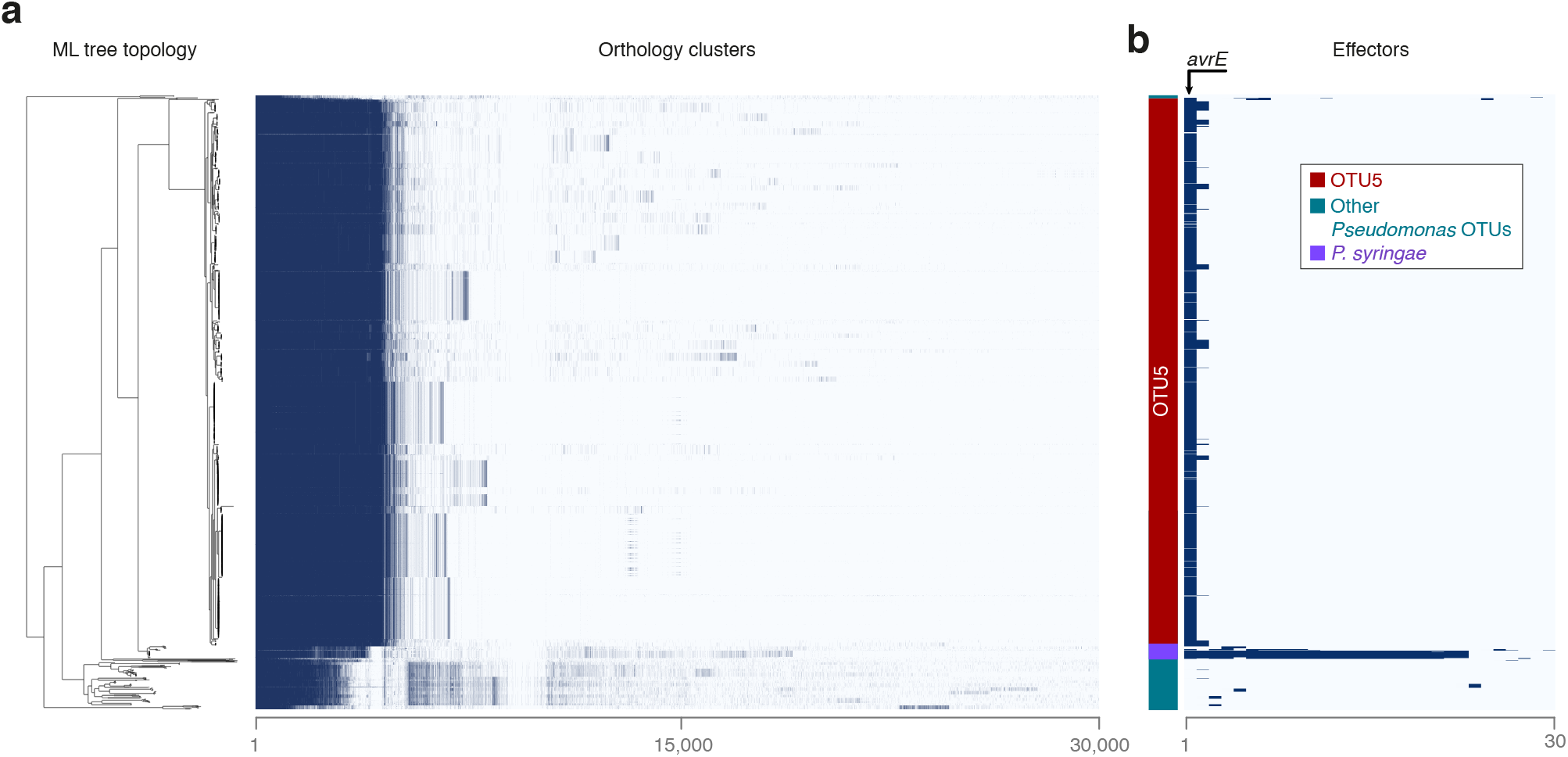
OTU5 strains vary in gene content but share an effector. (a) ML tree topology of 1,524 isolates and presence (dark blue) or absence (light blue) of the 30,000 most common orthologs as inferred with panX (Ding et al., 2018). OTU5 strains share 914 ortholog groups that are found at less than 10% frequency outside OTU5. (b) Presence/absence of 30 effector homologs. Only avrE homologs are present in more than 50 isolates. Related to Fig. S6.

To successfully colonize their hosts, microbes often deploy toxins, phytohormones and effectors that are secreted by the bacteria into host cells or the apoplast. To determine whether production of these compounds is likely to differ within OTU5 or between OTU5 and other clades, we generated a custom database of all effector genes and structural genes or enzymes for compounds known to be associated with host colonization in the genus *Pseudomonas.* We then used this database to independently annotate the effector as well as phytohormone and toxin biosynthesis gene content of each isolate (Fig. S6, Table S2).

OTU5 strains lacked all known genes for coronatine and syringomycin/syringopeptin synthesis, but auxin synthesis modules were found in almost all isolates (including isolates outside of OTU5). Genes for pectate lyase synthesis were broadly conserved both in and outside of OTU5 (Fig. S6). The *hrp-hrc* gene cluster, which encodes the type III secretion system (T3SS) along with effectors and several other proteins involved in pathogenicity (Alfano et al., 2000), is largely conserved across OTU5 isolates, with OTU5 alleles being most similar to the *hrp-hrc* clusters of previously sequenced *P. viridiflava* strains (Araki et al., 2006). We note that our search for plant-associated toxins and enzymes is not yet exhaustive. For example, other plant microbes deploy enzymes that can degrade the cell walls of their hosts (Almario et al., 2017), but such pathways have yet to be identified in *Pseudomonas.* Plants can detect microbes both through the presence of effector molecules and through microbe associated molecular patterns (MAMP). One such well-studied MAMP important in the *Pseudomonas-Arabidopsis* pathosystem is the flg22 peptide in flagellin (Gómez-Gómez et al., 1999). Isolates within OTU5 encode two major flg22 variants, which are highly divergent from one another (Table S2).

Effector proteins can increase bacterial fitness both through the suppression of the host immune system or through active promotion of proliferation in the plant (Chen et al., 2010; Xin et al., 2016) and are thought to be at the forefront of the coevolutionary interaction with the plant immune system (Karasov et al., 2014b). Only one gene for an effector homolog was broadly conserved across OTU5, *avrE.* It was shared with other *P. syringae* type isolates (Dillion et al., 2017), but found rarely outside this group. *avrE* encodes an effector that leads to increased humidity of the extracellular environment inside the plant, the apoplast (Xin et al., 2016). Experimental manipulation of apoplast humidity has shown that it is central to bacterial proliferation within the host. The most abundant *avrE* allele identified in our study is most similar to that previously observed in other *P. viridiflava* strains (Araki et al., 2006), with less similarity to the allele in the well-studied pathogen Pst DC3000 (Fig. S6). We thus conclude that the *avrE* homolog we identified in OTU5 is likely important for the success of OTU5 in the *A. thaliana* environment.

## Discussion

In the field of human microbiome and health, an understanding of how pathogen colonization differs between simplified clinical settings and more complex environments outside the clinic, which are often distinguished by levels of antibiotic treatment, has led to important innovations in disease treatment (Bakken et al., 2011). Understanding differences between pathogen colonization and evolution in natural versus agricultural systems may similarly lead to innovations that reduce pathogen pressure in agriculture (Hu et al., 2016). Much can be learned about the course of pathogen colonization and evolution by examining pathogen population diversity and demography (Hershberg et al., 2008; Yoshida et al., 2013). For example, whether pathogen expansions in host populations are composed of single, genetically monomorphic strains or instead comprise numerous genetically divergent strains can indicate whether the successful pathogen lineage is recently introduced/evolved, or has persisted over long time periods. Pathogen diversity is not only an indicator of the colonization process, but the diversity itself will also influence the course of colonization and the evolution of resistance in host populations (Karasov et al., 2014b).

In this study, we conducted a large-scale survey of *A. thaliana* leaves across populations and seasons to determine the most abundant OTUs of *Pseudomonas,* which includes important *A. thaliana* foliar pathogens. While we found a single OTU to be by far the most abundant *Pseudomonas* OTU across populations, this lineage, OTU5, is genetically diverse and consists of dozens, if not hundreds, of strains, diverged by approximately 300,000 years, with similar abilities to colonize the *A. thaliana* host. We were surprised to find a single dominant *Pseudomonas* lineage in the study area, given that wild *A. thaliana* populations can be colonized by a diversity of *Pseudomonas* pathogenic species (Jakob et al., 2002; Kniskern et al., 2011). We note, though, that while the OTU5 strains share many genetic features, they are not functionally synonymous--instead they are differentiated both at the level of gene content and the level of virulence. An important question for the future will be in how many other regions OTU5 is the dominant colonizer of *A. thaliana,* and how its genetic diversity is geographically structured across the entire host range.

The genetic diversity of OTU5 that we observed in this study stands in stark contrast to the monomorphic, recent pathogen spreads observed in typical agricultural epidemic systems (Cai et al., 2011; Wichmann et al., 2005). There are several non-mutually exclusive explanations for this. Industrial agricultural fields are often planted with one or a few plant genotypes, and environmental variation in these fields is reduced by fertilization prior to planting. The resulting uniformity of the field and host environment is known to influence the microbiota (Figuerola et al., 2015; Zhu et al., 2000), and to promote the expansions of single pathogens (Zhu et al., 2000). While we believe this to be the most likely explanation for the difference between our study system and agricultural settings, another possibility is that the diverse pathogenic expansions we observe in *A. thaliana* populations also occur in crop populations, but that such expansions may have gone unnoticed because their impact may be small in comparison to the monomorphic crop epidemics. Both theory (Leggett et al., 2013; Regoes et al., 2000) and observations (Ebert, 1998) have detailed scenarios in which specialized pathogens (such as those on crops) will proliferate to higher abundance in their hosts than will generalist pathogens.

Most studies of crop pathogen evolution have centered on the loss or gain of a single or a few virulence factors that subvert recognition by the host. Many instances of rapid turnover of virulence factors have been documented (Baltrus et al., 2011; Godfrey et al., 2011; Jackson et al., 2000), even within the span of a few dozen generations. In contrast, in the *A. thaliana* system, we observe long-term stability of the *avrE* effector gene. The long-term success of OTU5 suggests that genetic factors leading to the success of divergent strains are likely to be conserved across these strains. Beyond the molecular mechanism of avrE-dependent virulence, a growing number of studies has demonstrated that *avrE* may be central to the success of several plant pathogens and on several plant hosts. *avrE* homologs have not only been found in *Pseudomonas,* but have also been identified in other bacterial taxa, where they have been implicated in pathogenicity as well. DspE, an AvrE homolog in the plant pathogen *Erwinia amylovora,* functions similarly to AvrE (Bogdanove et al., 1998), pointing to many pathogens relying on the AvrE mechanism to enhance their fitness. Hosts often have evolved means to detect effector proteins (Chisholm et al., 2006). While several soybean cultivars can recognize the activity of AvrE (Kobayashi et al., 1989), gene-for-gene resistance to the avrE-containing Pst DC3000 model pathogen has so far not been found in *A. thaliana* (Velásquez et al., 2017) nor has quantitative resistance been observed, even though the effector reliably enhances colonization of Pst DC3000 (Xin et al., 2016).

It is reasonable to hypothesize that the plant host has evolved mechanisms that suppress the disease effect of OTU5. By itself, many OTU5 strains can reduce plant growth in gnotobiotic culture by more than 50% or even kill the plant. In natural populations, the pathogenic effect appears to be mitigated, since we isolated OTU5 strains from plants that did not appear to be heavily diseased. Indeed, several environmental and genetic factors are known to affect the the pathogenic effect of microbes including the physiological state of the plant (MacQueen and Bergelson, 2016) and the presence of other microbiota (Goss and Bergelson, 2007; Innerebner et al., 2011; Mendes et al., 2011). Understanding mechanisms of disease-mitigation in response to OTU5 will provide insight into how natural plant populations can blunt the effects of a common pathogen without instigating an arms race, and thereby suggest possible novel approaches to disease-protection in agriculture.

## Experimental Procedures

### Sample collection

For the 16S rDNA survey, *A. thaliana* samples were collected from five to six populations (sites) around Tübingen (Fig. S1 a), in the fall and spring of 2014, 2015 and 2016; the number of sampled plants is indicated in Fig. 1a. For endophytic and epiphytic sample fractionation, whole rosettes were processed as described in ref. (Agler et al., 2016). Briefly, rosettes were washed once in water for 30 s, then in 3-5 mL of epiphyte wash solution (0.1% Triton X-100 in 1x TE buffer) for 1 min, before filtering the solution through a 0.2 μm nitrocellulose membrane filter (Whatman, Piscataway, NJ, USA) to collect the epiphytic fraction. For the endophytic fraction, the initial rosette was surface sterilized by washing with 80% ethanol for 15 seconds followed by 2% bleach (sodium hypochlorite) for 30 seconds, before rinsing three times with sterile autoclaved water. Samples were stored in screw cap tubes and directly frozen in dry ice. DNA extraction was conducted following (Agler et al., 2016), including a manual sample grinding step followed by a lysis step with SDS, Lysozyme and proteinase K, a DNA extraction step based on phenol-chloroform and a final DNA precipitation step with 100% ethanol.

Additional samples were collected from two of the six sites sampled for 16S rDNA, from Eyach, on December 11, 2015, and March 23, 2016, and from Kirchentellinsfurt on December 15, 2016, and March 31, 2016. Whole rosettes were removed with sterile scissors and tweezers, and washed with deionized water. Two leaves were removed and independently processed, and the remaining rosette was flash-frozen on dry ice. The flash-frozen material was processed for metagenomic sequencing and 16S rDNA sequencing of the v4 region. The removed leaves were placed on ice, washed in 70%-80% EtOH for 3-5 seconds to remove lightly-associated epiphytes. Sterilized plants were ground in 10 mM MgSO**4** and plated on King's Broth (KB) plates containing 100 μg/mL nitrofurantoin (Sigma). Plates were incubated at 25°C for two days, then placed at 4°C. Colonies were picked randomly from plates between 3-10 days after plating, grown in KB with nitrofurantoin overnight, then stored at-80°C in 15-30% glycerol.

### 16S v3-v4 amplicon sequencing

The 16S v3-v4 region was amplified as described (Agler et al., 2016). Briefly, PCR reactions were conducted using a two-step protocol using blocking primers to decrease plant plastid 16S rDNA amplification. The first PCR was conducted with primers B341F / B806R in 20 μL reactions containing 0.2 μL Q5 high-fidelity DNA polymerase (New England Biolabs, Ipswich, MA, USA), 1x Q5 GC Buffer, 1x Q5 5x reaction buffer, 0.08 μM each of forward and reverse primer, 0.25 μM blocking primer and 225 μM dNTP. Template DNA was diluted l:l in nuclease free water and 1 μL was added to the PCR. Triplicates were run in parallel on three independent thermocyclers (Bio-Rad Laboratories, Hercules, CA, USA); cycling conditions were 95 °C for 40 s, 10 cycles of 95 °C for 35 s, 55 °C for 45 s, 72 °C for 15 s, and a final elongation at 72 °C for 3 min. The three reactions were combined and 10 μL were used for enzymatic cleanup with Antarctic phosphatase and Exonuclease I (New England Biolabs; 0.5 μL of each enzyme with 1.22 μL Antarctic phosphatase buffer at 37°C for 30 minutes followed by 80°C for 15 min). One microliter of cleaned PCR product was subsequently used in the second PCR with tagged primers including the Illumina adapters, in 50 μL containing 0.5 μL Q5 high-fidelity DNA polymerase (New England Biolabs), 1x Q5 GC Buffer, 1x Q5 5x reaction buffer, 0.16 μM each of forward and reverse primer and 200 μM dNTP. Cycling conditions were the same as for the first PCR except amplification was limited to 25 cycles. The final PCR products were cleaned using 1.8x volume Ampure XP purification beads (Beckman-Coulter, Brea, CA, USA) and eluted in 40 μL according to manufacturer instructions. Amplicons were quantified in duplicates with the PicoGreen system (Life Technologies, Carlsbad, CA, USA) and samples were combined in equimolar amounts into one library. The final libraries were cleaned with 0.8x volume Ampure XP purification beads and eluted into 40 μL. Libraries were prepared with the MiSeq Reagent Kit v3 for 2x300 bp paired-end reads (Illumina, San Diego, CA, USA). All the samples were analysed in 9 runs on the same Illumina MiSeq instrument. Samples failing to produce enough reads on one run were re-sequenced and data from both runs were merged. The raw sequencing data was deposited at the National Center for Biotechnology Information (NCBI) Short Read Archive under BioProject PRJNA430505.

### 16S rDNA v3-v4 amplicon data analysis

Amplicon data analysis was conducted in Mothur (Schloss et al., 2009). Paired-end reads were assembled *(make.contigs)* and reads with fewer than 5 bp overlap (full match) between the forward and reverse reads were discarded *(screen.seqs).* Reads were demultiplexed, filtered to a maximum of two mismatches with the tag sequence and a minimum of 100 bp in length. Chimeras were identified using Uchime in Mothur with more abundant sequences as reference *(chimera.uchime,* abskew=1.9). Sequences were clustered into OTUs at the 99 % similarity threshold using VSEARCH in Mothur with the distance based clustering method (dgc) *(cluster).* Individual sequences were taxonomically classified using the rdp classifier method *(classify.seqs,* consensus confidence threshold set to 80) and the greengenes 16S rDNA database (13_8 release). Each OTU was taxonomically classified *(classify.otu,* consensus confidence threshold set to 66), non-bacterial OTUs and OTUs with unknown taxonomy at the kingdom level were removed, as were low abundance OTUs (< 50 reads, *split.abund).* The confidence of OTU classification to the genus *Pseudomonas* was at least 97%. The most abundant sequence within each *Pseudomonas* OTU was selected as the OTU representative for phylogenetic analyses.

All statistical analyses were conducted in R 3.2.3 (Team and Others, 2010). In order to avoid zero values, relative abundance data was transformed using a log (x+a) formula where *a* is the minimum value of the variable divided by two. Normality after transformation was assessed using Shapiro Wilk's normality test. Factors influencing *Pseudomonas* relative abundance were studied using multi-factorial ANOVA. When necessary, sites PFN and K6911 were excluded from the analysis, as they had missing data points (Fig. 1a). Mean differences were further verified with Wilcoxon's non-parametric test. Differences between *Pseudomonas* populations were assessed by calculating Bray-Curtis dissimilarities between samples using the “vegdist” function of the vegan package(Oksanen et al., 2007). These distances were used for principal coordinates analysis using the “dudi.pco” function of the ADE4 package(Dray et al., 2007), and for PERMANOVA to study the effect of different factors on the structure of *Pseudomonas* populations using the “Adonis” function of the vegan package.\

The 16S rDNA analysis of 192 plants in Eyach for which also metagenomic shotgun data were generated (see below) involved the amplification of the v4 region using the published primers 515F-806R (Schmidt et al., 1991) on an Illumina Miseq instrument with 2x250 bp paired end reads, which were subsequently merged. Sequences were clustered with uclust (Edgar, 2010), 99% identity, and taxonomically assigned using the RDP taxonomical assignation (Wang et al., 2007). 103 OTUs were assigned to the genus *Pseudomonas.* One OTU aligned with 100% identity over its entire length to OTU5, the most abundant OTU identified in the cross-population survey of the v3-v4 region described above.

### Metagenomic assessment of bacterial load

Total DNA was extracted from flash-frozen rosettes by pre-grinding the frozen plant material to a powder using a mortar and pestle lined with sterile (autoclaved) aluminum foil and liquid nitrogen as needed to keep the sample frozen. Between 100 mg and 200 mg of plant material were then transferred with a sterile spatula to a 2 mL screw cap tube (Sarstedt) containing 0.5 mL of 1 mM garnet rocks (BioSpec). To this, 800 |L of room temperature extraction buffer was added, containing 10 mM Tris pH 8.0, 10 mM EDTA, 100 mM NaCl, and l.5% SDS. Lysis was performed in a FastPrep homogenizer at speed 6.0 for 1 minute. These tubes were spun at 20,000 x g for 5 minutes, and the supernatant was mixed with *Ys* volume of 5M KOAc in new tubes to precipitate the SDS. This precipitate was in turn spun at 20,000 x g for 5 minutes and DNA was purified from the resulting supernatant using Solid Phase Reversible Immobilisation (SPRI) beads (DeAngelis et al., 1995) at a bead to sample ratio of 1:2. DNA was quantified by PicoGreen, and libraries were constructed using a Nextera protocol modified to include smaller volumes (similar to (Baym et al., 2015)). Library molecules were size selected on a Blue Pippin instrument (Sage Science, Beverly, MA, USA). Multiplexed libraries were sequenced with 2x150 bp paired-end reads on an HiSeq3000 instrument (Illumina).

A significant challenge in the analysis of plant metagenomic sequences, is the proper removal of the host DNA. In order to remove host derived sequences, reads were mapped against the *A. thaliana* TAIR10 reference genome with bwa mem(Li, 2013) using standard parameters. Subsequently, all read pairs flagged as unmapped were isolated from the main sequencing library with samtools (Li et al., 2009) as this represents the putatively “metagenomic” fraction.

Afterwards, this metagenomic fraction was mapped against the NCBI nr database (NCBI Resource Coordinators. Database resources of the National Center for Biotechnology Information 2016) with the blastx implementation of DIAMOND (Buchfink et al., 2015) using standard parameters.

Based on the reference sequences for which our metagenomic reads had significant alignments, taxonomic binning of sequencing data was performed with MEGAN via the naive LCA algorithm (Huson et al., 2007). Normalization of binned reads was performed with custom scripts and based on the number of reads binned into any given genus including reads assigned to species in that genus, taxa abundance was estimated. Metagenomic short read sequences were deposited in the European Nucleotide Archive (ENA) under the Primary Accession PRJEB24450.

### Whole-genome sequencing

Bacterial DNA, both genomic and plasmid, was extracted using the Puregene DNA extraction kit (Invitrogen). Single bacterial colonies were grown overnight in Luria broth+100 μg/mL Nitrofurantoin in 96-well plates. Plates were spun down for 10 minutes at 8000g, then the standard Puregene extraction protocol was followed. The capacity of the protocol to extract plasmid DNA was verified by extracting the DNA from a strain whose plasmids were previously identified (Pst DC3000)(Buell et al., 2003). Primers specific to these plasmids successfully amplified the puregene-extracted sample.

Genomic and plasmid DNA libraries for single bacteria and for whole plant metagenomes were constructed using a modified version of the Nextera protocol (Caruccio, 2011), modified to include smaller volumes (Rowan et al 2017). Briefly, 0.25-2ng of extracted DNA was sheared with the Nextera Tn5 transpososome. Sheared DNA was amplified with custom primers for 14 cycles. Libraries were pooled and size-selected for the 300-700bp range on a Blue Pippin. Resulting libraries were then sequenced on the Illumina HiSeq3000. Coverage and assembly statistics are detailed in Fig. S2.

### Assembly and annotation

Genomes were assembled using Spades (Bankevich et al., 2012) (standard parameters) and assembly errors corrected using pilon (Walker et al., 2014) (standard parameters). Gene annotations were achieved using Prokka (Seemann, 2014) (standard parameters). Those genomes with N50<25kbp or less than 3000 annotated genes were deemed to be of insufficient quality and were excluded from further analyses except for 19 genomes sequenced in the second season. Distributions of gene number and assembly quality are displayed in Fig. S3. The number of missing genes per genome was assessed using Busco (Simão et al., 2015). Assembled genomes were deposited in the European Nucleotide Archive (ENA) under the Primary Accession PRJEB24450.

Because Prokka does not successfully identify several effectors, in addition to other genes involved in interactions with the host, we augmented the Prokka annotation with several additional annotation sets. We predicted genes on the raw genome FASTA sequences using AUGUSTUS-3.3 (Stanke and Waack, 2003) and-genemodel=partial-gff3=on-species=E_coli_K12 settings. The protein sequence of each predicted gene was extracted using a custom script.

We annotated effectors using BLASTP-2.2.31+ (Altschul et al., 1990) specifying the AUGUSTUS predicted proteomes as query input and the Hop database (http://www.Pseudomonas-syringae.org/T3SS-Hops.xls) as reference database. We filtered the BLASTP results with a 40% identity query to reference sequence threshold, a 60% alignment length threshold of query to reference sequence and a 60% length ratio threshold of query and reference sequence (empirically determined). Hits of interest were manually extracted and controlled using online BLASTP and NCBI conserved domain search.

Toxins and phytohormones were annotated using the same BLASTP settings as described for effectors. We used custom NCBI protein databases including a set of genes involved in the toxin synthesis pathway. A strain was scored as toxin pathway encoding if all selected components of a pathway were present. Hrp-hrc clusters were also annotated using the formerly described BLAST and filtering settings and *P. syringae pv tomato DC3000* and *P. viridiflava PNA 3.3* as reference sequences.

### Pan-genome analysis and phylogenetics

The panX pan-genome pipeline was used to assign orthology clusters (Ding et al., 2018) and build alignments of these clusters that were then used for phylogenetic analysis in RAxML (Stamatakis et al., 2005). The parameters used were the following: divide-and-conquer algorithm (-dmdc) was used on the diamond clustering, a subset size of 50 was used in the dmdc (-dcs 50), a core genome cutoff of 70% (-cg 0.7).

Whole-genome phylogenies of the strains were constructed using RAxML (Stamatakis et al., 2005) using the gamma model of rate heterogeneity and the generalized time reversible model of substitution. The phylogenies were built from all SNPS present in the concatenated core genomes of strains identified by panX. Eight-hundred and seven genes were considered as core. We performed 100 bootstrap replicates in RAxML to establish the confidence in the full tree.

Within the 1355 isolates belonging to OTU5, 107 distinct strains were represented. One representative of each strain was picked at random, then recombination importation events were identified among these 107 strains using ClonalFrameML (Didelot and Wilson, 2015). ClonalFrameML estimated a high recombination rate within OTU5, estimating that a substitution in the tree was six times more likely to result from a recombination event than a mutation event. Specifically, ClonalFrameML estimated the following parameters: the 1/δ parameter (inverse importation event tract length in bp) was estimated as 7.79x10^-3^/bp (var=2.18014^-9^) and the Posterior Mean ratio between the probability of recombination (R) and the nucleotide diversity, θ, was R/θ =1.19 (var=5.07x10^-5^). The estimated sequence divergence between imported tracts and the acceptor genome v=0.04 (var=l.13x10^-8^). The relative effect of recombination over mutation r/m=(R/θ) x V x δ = 6.18. Recombination tracts were removed from the alignments, and the remaining putatively non-recombined strict core genes (present in all 107 genomes) were used for subsequent dating of coalescence.

To estimate the age of OTU5 we considered only those ortholog groups that were conserved across all 107 OTU5 strains. These orthologs were concatenated and ClonalFrameML (Didelot and Wilson, 2015) was used to identify recombination tracts that could inflate the branch length of members of the OTU as described above. TMRCA of the OTU was estimated by calculating the mid-point-root to tip sequence divergence for a representative of all 107 strains within OTU5, then dividing the median value of this distance by the neutral substitution rate (Kimura, 1968) (we used here the point estimate of 8.7x10^-8^ with our estimate of sd=6.0x10^-8^ (McCann et al., 2017)). While we consider all sites (degenerate and non-degenerate) in the putatively non-recombined core, in addition to the fact that substitution rate is likely inaccurate for the longer timescale analysed in the present study, both of these inaccuracies would likely lead to the underestimation of the age.

### Pathogenicity assays

The plant genotype Eyach 15-2 (CS76399), collected from Eyach, Germany, was previously determined to represent a plant genetic background common to the geographical region. Seeds were sterilized by overnight incubation at-80°C, then 4 hours of bleach treatment at room temperature (seeds in open 2 ml tube in a desiccator containing a beaker with 40 ml Chlorox and 1 ml HCl (32%)). The seeds were then stratified for three days at 4°C in the dark on ½ MS media. Plants were grown in 3-4 mL ½ MS medium in six-well plates in long-day (16 hours) at 16°C. 12-14 days after stratification, plants were infected with single bacterial strains.

Bacteria were grown overnight in Luria broth and the relevant antibiotic (either 10 μ/mL of Kanamycin or Nitrofurantoin), diluted 1:10 in the morning and grown for 2 additional hours until they entered log phase. The bacteria were pelleted at 3500 g, resuspended in 10 mM MgSO**4** to a concentration of OD**600**=0.01. 200 μl of bacteria were drip-inoculated with a pipette onto the whole rosette. Plates were sealed with parafilm and returned to the growth percival. Seven days after infection, whole rosettes were cut from the plant and fresh mass was assessed.

For growth assays of dead bacteria, we performed growth and dilution of bacteria as above, then boiled the final preparation at 95°C for 38 minutes. Plants were treated with the dead bacteria in the same manner as described above.

## Acknowledgements

We thank Marek Kucka for methodological help with Tn5 transpososome purification, Julia Vorholt for recommendations on infection methods, Matthew Clark for providing protocols for amended Nextera methodology, Hernán Burbano for discussions, and Joy Bergelson, Jeff Dangl and Michael Werner for critical reading of the manuscript. Funding was provided by HFSP long-term fellowships (TLK, DSL), an EMBO LRTF (TLK), ERC AdG IMMUNEMESIS (DW) and the Max Planck Society (DW, EK).

## Author Contributions

TLK, EK and DW devised the study. TLK, JA, CF, MG, SK, DSL, and MN performed the experiments, TLK, JA, WD, DSL, MG, JR, and RAN analyzed the data. DH advised on library preparation methods. RAN, EK and DW advised on data analysis. TLK, JA and DW wrote the manuscript with help from all authors.

## Supplementary Information

Table S1. Collection Locations and dates for all samples.

Table S2. Genes annotated in plant-associated pathways and amino acid sequence of flg22 variants in OTU5.

Fig. S1. Changes in *Pseudomonas* populations colonizing *A. thaliana* leaves.

Fig. S2. *Pseudomonas* OTUs and whole-genome phylogeny.

Fig. S3. *De novo* genome assembly of 1,524 strains sequenced in this study.

Fig. S4. Leaf-endophytic *Pseudomonas* diversity at Eyach site. overlap between amplicon sequencing and strain isolation data.

Fig. S5. Gnotobiotic trial with OTU5 strains.

Fig. S6. Toxin and phytohormone distribution.

**Table S1.** Collection Locations and dates for all samples.

| Site | Latitude | Longitude | Sample type |
| --- | --- | --- | --- |
| EY | 48.446111 | 8.781611 | 16S, metagenome, isolates |
| K6911 | 48.541278 | 9.0925 | 16S |
| PFN | 48.561087 | 9.109294 | 16S |
| JUG | 48.555722/48.556255 | 9.134833/9.135424 | 16S, metagenome, isolates |
| WH | 48.506827 | 8.936418 | 16S |
| ERG | 48.495362 | 8.809083 | 16S |

**Table S2.**
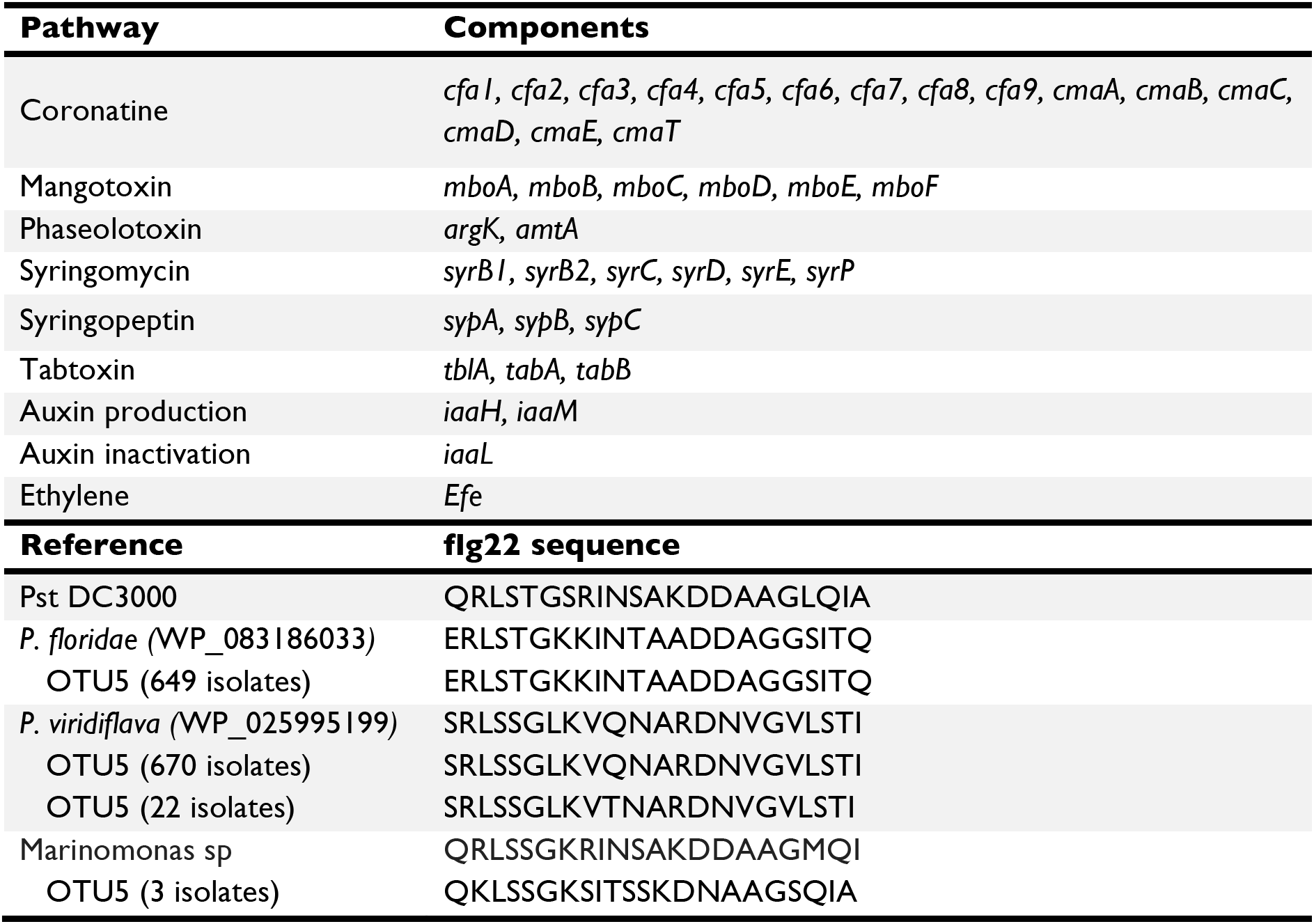
Genes annotated in plant-associated pathways and amino acid sequence of flg22 variants in OTU5. Top. The listed genes (right column) were custom-annotated in each of the genomes to ascertain the presence of the listed pathway (left column). Bottom. Left column lists the genome from which flg22 sequence was extracted. OTU5 encodes two major variants of flg22, one similar to a *P. floridae* genotype, and the other to a *P. viridiflava* genotype.

**Fig. S1.**
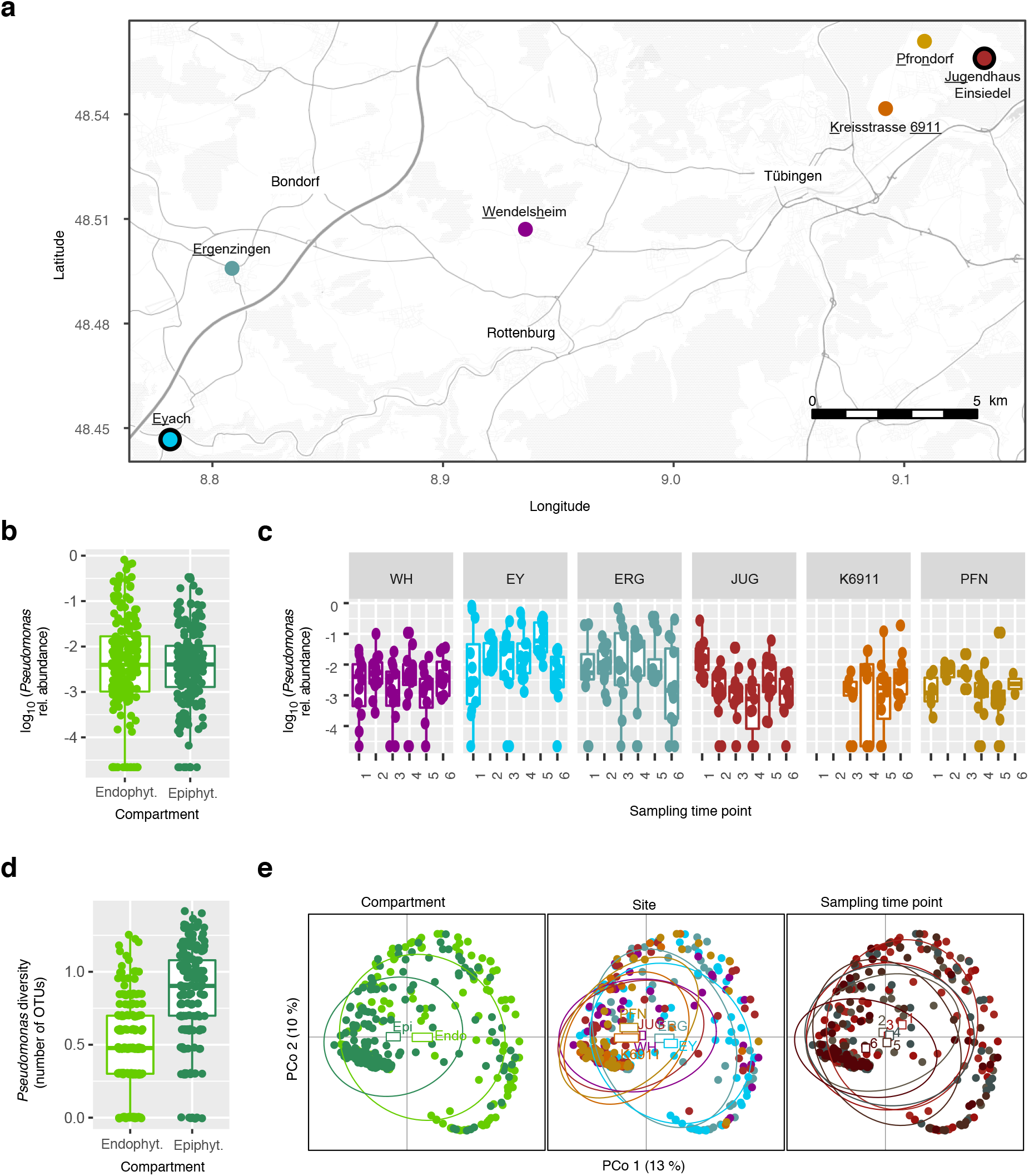
Changes in *Pseudomonas* populations colonizing *A. thaliana* leaves. (a) Location of the sampling sites around Tübingen (Germany). The two sites from which isolates were cultured indicated by black outlines. (b) *Pseudomonas* abundance in the endophytic (Endo.) and epiphytic (Epi.) compartments. (c) *Pseudomonas* abundance across the different sites and sampling time points (see Fig. 1a). (d) *Pseudomonas* diversity in the endophytic (Endo.) and epiphytic (Epi.) compartments. (e) Principal coordinates analysis (PCoA) based on Bray-Curtis distances, depicting the differences between *Pseudomonas* populations across the different compartments, sites and sampling time points. *Pseudomonas* relative abundance (RA) was calculated as the ratio of *Pseudomonas* reads to the total number of bacterial reads. Related to Fig. 1.

**Fig. S2.**
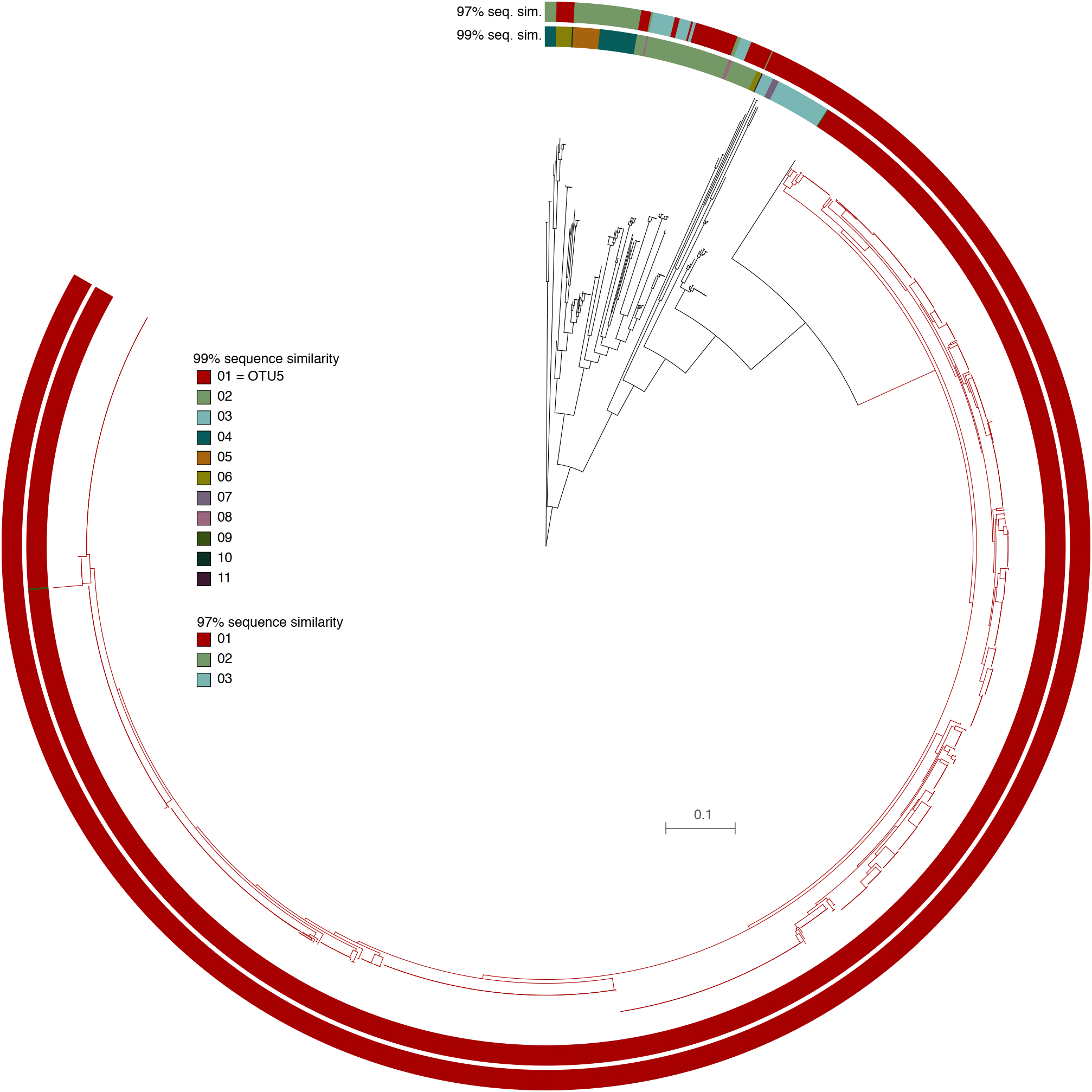
*Pseudomonas* OTUs and whole-genome phylogeny. 16S rDNA sequences were extracted from 1,524 *Pseudomonas* isolates with whole-genome sequences. Clustering based on 99% or 97% sequence similarity of the v3-v4 region of the 16S rDNA is compared to an ML core genome phylogenetic tree. Colors indicate group, ranked by abundance. The most abundant group corresponds to OTU5 (Bordeaux color), with clustering at 99% sequence identity being more consistent with the core genome tree than 97% clustering. Related to Fig. 1, 2 and 3.

**Fig. S3.**
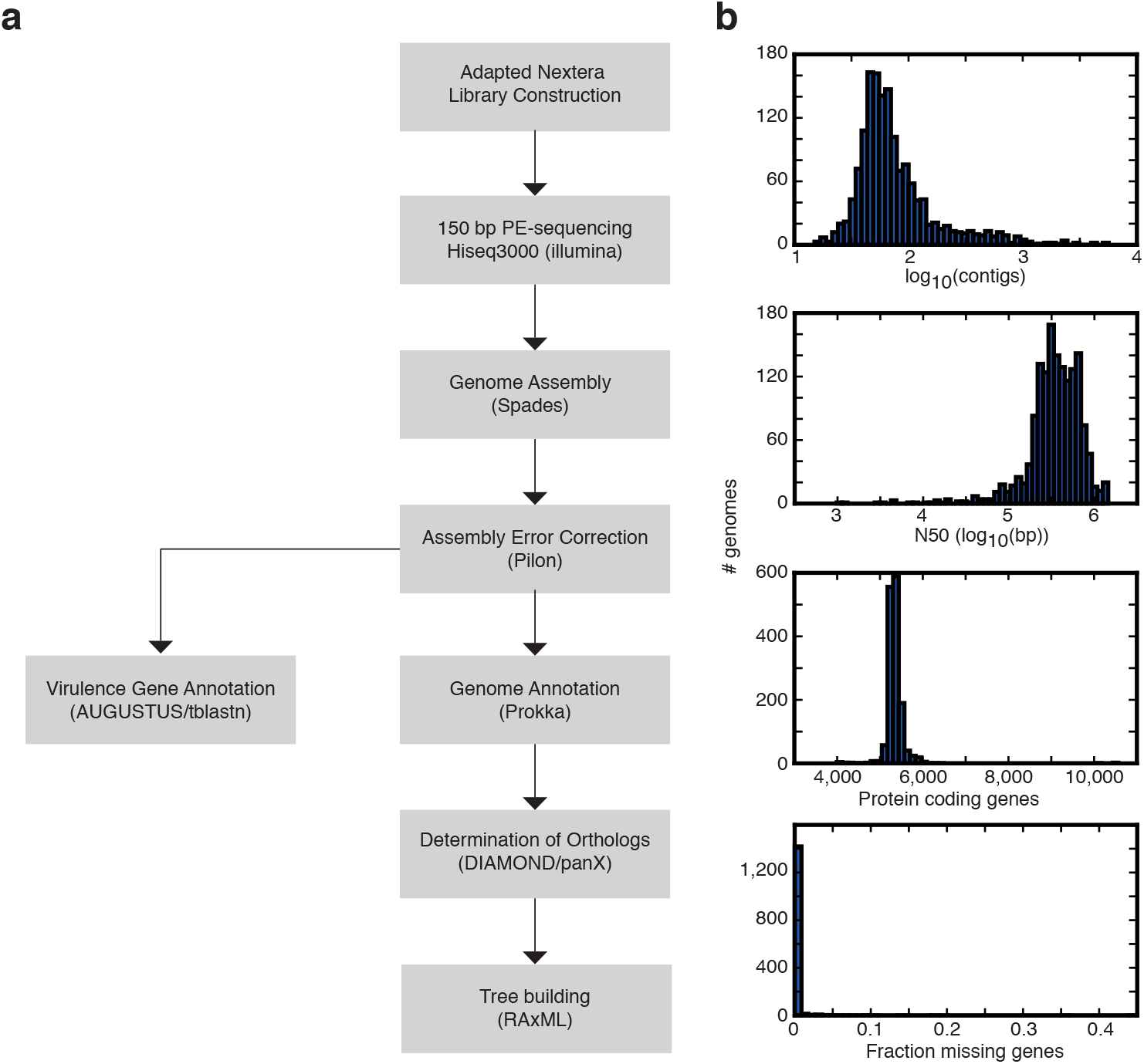
*De novo* genome assembly of 1,524 strains sequenced in this study. Genomes were filtered for basic assembly quality (N50>25,000bp, and more than 3500 genes per genome). The number of contigs in the assembled genome (a) the N50 for each assembly (b) the number of protein coding genes annotated per genome (c) and the percentage of genes predicted to be missing per assembly (d) are shown for the 1,524 genomes remaining after filtering. Related to Fig. 4.

**Fig. S4.**
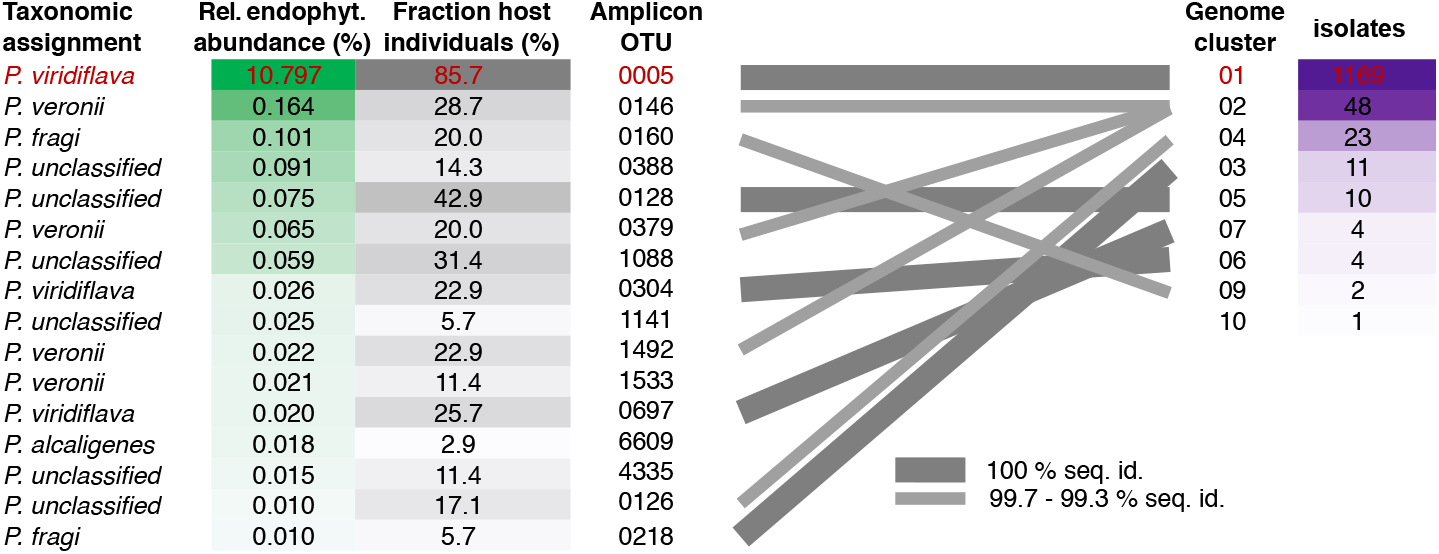
Leaf-endophytic *Pseudomonas* diversity at Eyach site: overlap between amplicon sequencing and strain isolation data. Memberships in strain clusters identified by 99% sequence similarity of the 16S rDNA v3-v4, either through amplicon sequencing (OTUs, left) or strain isolation (genomes, right). Matching groups (grey lines) were identified by comparing representative sequences for the amplicon OTUs/genome clusters. Even when considering data from different host individuals sampled at different times there is a good overlap between the two collections. Related to Fig. 5.

**Fig. S5.**
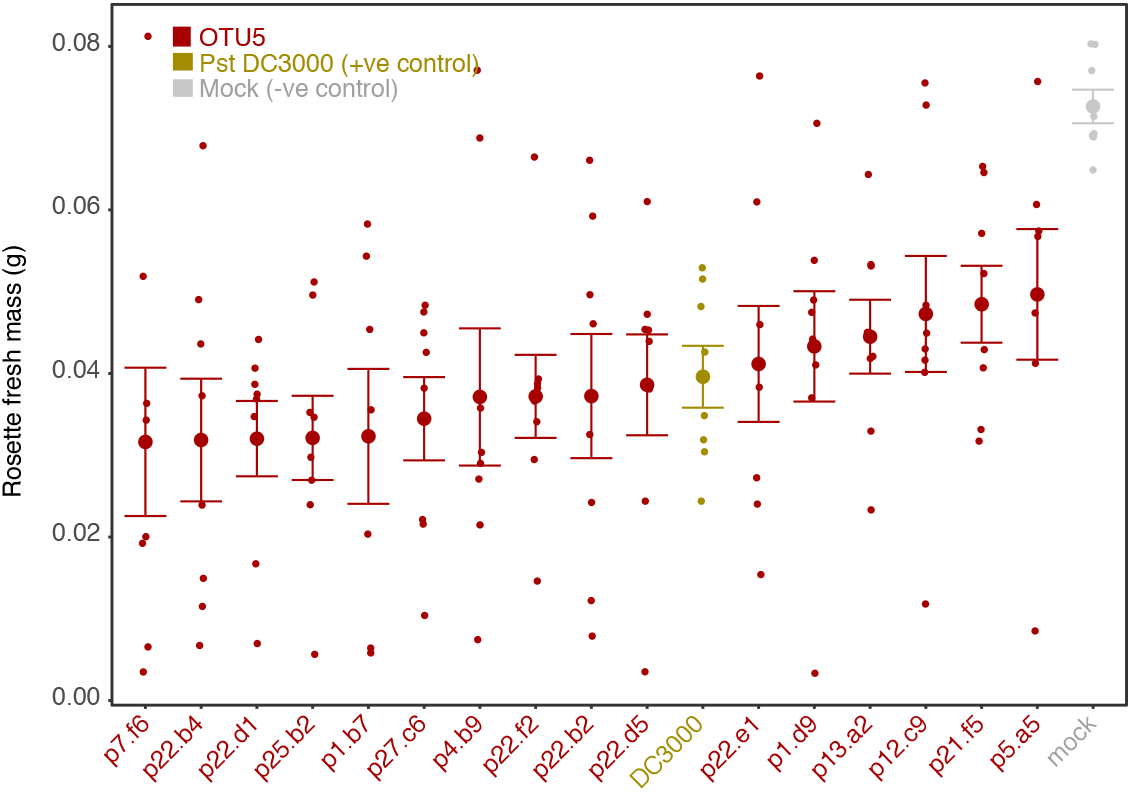
Gnotobiotic trial with OTU5 strains. 14 day old *A. thaliana* plants of accession Eyach 15-2 were drip-infected with different OTU5 strains. All tested strains significantly reduced plant growth (n=8 replicates per sample, Student's t-test, q-value<0.05). Related to Fig. 3c.

**Fig. S6.**
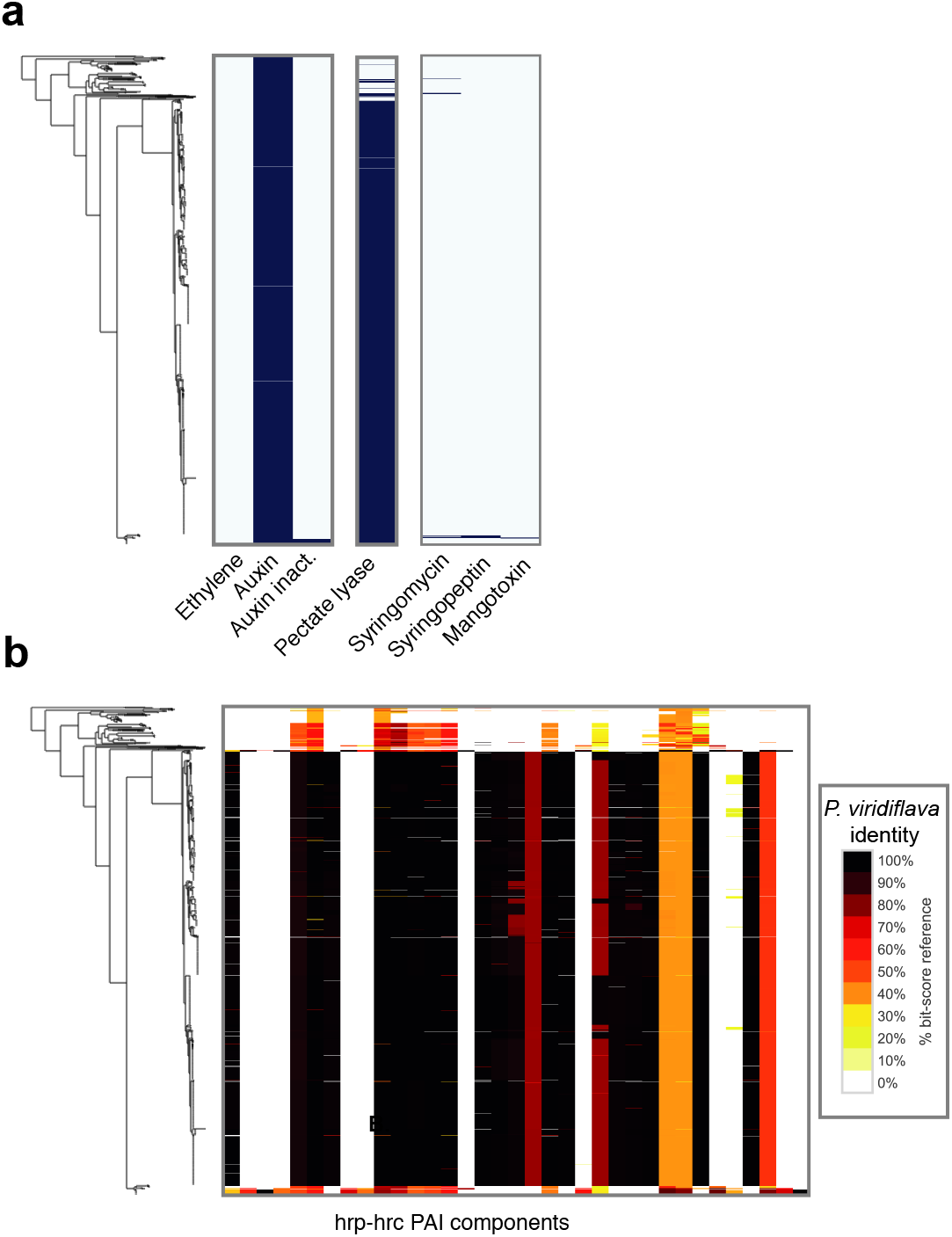
Toxin and phytohormone distribution. Toxins and phytohormones were annotated in the 1,524 genomes with a custom database and the genetic elements described in Table S2. Related to Fig. 6.

